## Supplementary figures for "Agent-Based Modeling of T Cell Receptor Cooperativity"

**Supplementary Materials**

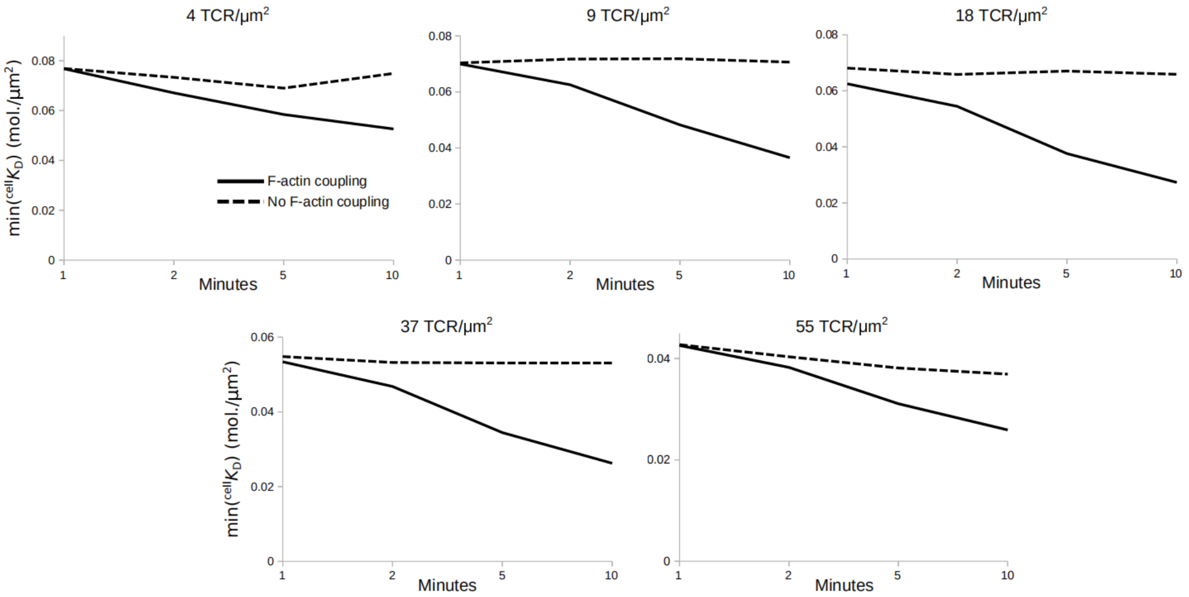


**Figure S1.** Time evolution of min($^{cell}K_D$) with TCR and pMHC titration, in the presence (solid lines) or absence (dashed lines) of F-actin centripetal transport.

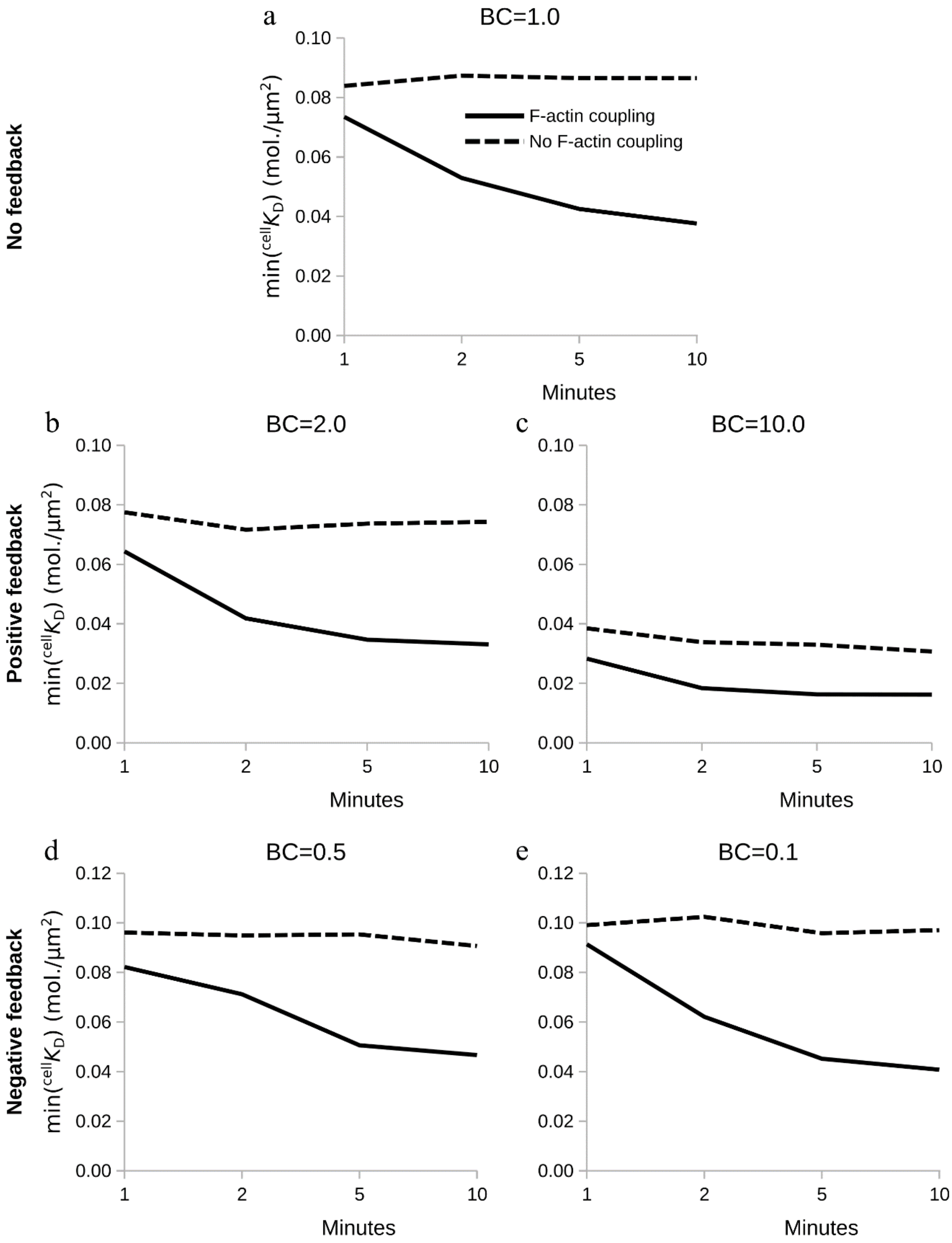


**Figure S2.** Time evolution of min($^{cell}K_D$) with feedback from the binding coefficient, *B*, in the presence or absence of centripetal transport of complexes.
